# High-throughput screening and selection of PCB-bioelectrocholeaching, electrogenic microbial communities using single chamber microbial fuel cells based on 96-well plate array

**DOI:** 10.1101/2021.06.09.447729

**Authors:** L. Szydlowski, J. Ehlich, N. Shibata, I. Goryanin

**Affiliations:** Okinawa Institute of Science and Technology, Malopolska Centre of Biotechnology Jagiellonian University; Brno University of Technology; Okinawa Institute of Science and Technology; Okinawa Institute of Science and Technology, University of Edinburgh, Tianjin Insititute of Biotechnology

## Abstract

We demonstrate a single chamber, 96-well plate based Microbial Fuel Cell (MFC). This invention is aimed at robust selection of electrogenic microbial community under specific conditions, (pH, external resistance, inoculum) that can be altered within the 96 well plate array. Using this device, we selected and multiplicated electrogenic microbial communities fed with acetate and lactate that can operate under different pH and produce current densities up to 19.4 A/m^3^ (0.6 A/m^2^) within 5 days past inoculation. Moreover, studies shown that Cu mobilization through PCB bioleaching occurred, thus each community was able to withstand presence of Cu^2+^ ions up to 600 mg/L. Metagenome analysis reveals high abundance of *Dietzia* spp., previously characterized in MFCs, but not reported to grow at pH 4, as well as novel species, closely related to *Actinotalea ferrariae*, not yet associated with electrogenicity. Microscopic observations (combined SEM and EDS) reveal that some of the species present in the anodic biofilm were adsorbing copper on their surface, probably due to the presence of metalloprotein complexes on their outer membranes. Taxonomy analysis indicated that similar consortia populate anodes, cathodes and OCP controls, although total abundances of aforementioned species are different among those groups. Annotated metagenomes showed high presence of multicopper oxidases and Cu-resistance genes, as well as genes encoding aliphatic and aromatic hydrocarbon-degrading enzymes. Comparison between annotated and binned metagenomes from pH 4 and 7 anodes, as well as their OCP controls revealed unique genes present in all of them, with majority of unique genes present in pH 7 anode, where novel *Actinotalea* spp. was present.

## 1. Introduction

Microbial Fuel Cells (MFCs) are a type of chemical fuel cell where the anodic reaction is catalysed by various microorganisms oxidizing organic matter. When coupled to cathodic reduction of oxygen, this system yields energy in the form of electricity. Given the exponential growth of studies focused on MFCs and electrogenic bacteria in general [1], numerous reactor designs have been demonstrated. However, singularities of those systems renders reproducibility of experiments extremely difficult.

Prior focus on unifying reactor conditions to study EET was on manufacturing multiple stand-alone microbial reactors that could be manufactured e.g. based on small glass vials[2,3]. Reactors design, materials used, dimensions and electrochemical properties (e.g. internal resistance) differed between each research group. Hou et al.[4–6] demonstrated the use of 24 well plate array with microfabricated gold electrodes with ferricyanide [4], air-cathodes [5] or microfluidic channels with continuous anolyte and catholyte replenishment[6], which increased the power output by a factor of three. These reactors were used to screen previously selected electrochemically active environmental isolates. Another example of well-plate array implementation was demonstrated by Yuan *et al*., ([7]), where extracellular electron transfer was coupled to the color change of the probe. Recently, Molderez *et al*. [8] constructed a 128-channel potentiostat connected to a printed circuit board (PCB) microarray. The entire microarray was immersed in an anolyte solution and supplied with a reference electrode to perform a high-throughput investigation pertaining to the effect of the anodic potential on electroactive biofilm growth. Zhou *et al*. [9] proposed a well-plate-based, high-throughput colorimetric assay for microbial electrochemical respiration, which would indicate extracellular electron transfer. Alternatively, a 48-well plate with hydrophobic wax layer separating electrodes was developed by Choi *et al*. [10], followed up by single sheet paper-based electrufluidic array (8 cells)[11]). Later, Tahernia and colleagues [12] developed paper-based, disposable 64-well array yielding power densities up to 23 μW/cm^2^. The device has been successfully used to characterize electrochemical properties of various *Shewanella oneidensis* and *Pseudomonas aeruginosa* mutants. Eventually, 96-well electrofluidic array was developed using the same fabrication method[13].

Nonetheless, the aforementioned systems [10–12,14,15] have low internal volume, which may not be suitable for experiments requiring continuous operation in order of multiple days or weeks. Moreover, the aforementioned wax paper designs are disposable and thus new device needs to be manufactured for every study.

In this paper, we present miniaturized MFCs based on 96 well plate array, which is reusable and durable for studies that took. Using our device, we screened for electrogenic consortia derived from two different inocula (air conditioning outflow and mangrove sediments) fed with identical substrate. We also applied different pH conditions to those consortia within identical reactors to allow multiparameter, comparative studies in their electrochemical performance. Moreover, we sequenced anodic biofilms from all tested groups and visualized them under electron microscopy.

## 2. Methods

### 2.1. MFC Construction

Each well was built as an individual MFC in a membraneless design (Fig.1). Base anode plate was built of a standard Printed Circuit Board (PCB) with a thin copper sheet (17 μm) electrolessly plated with nickel and covered with a thin layer of immersion gold (ENIG – RoHS by JLCPCB China). This part was exposed to the anolyte solution inside of the reactor. Volume of each well was 0.577 cm^3^ and spacing between electrodes was 1 cm. Surface area of the anode was enhanced by placing carbon veil discs on top of the base metal plate. Base cathode material was carbon paper (090, Toray, Japan) treated with polytetrafluoroethylene to prevent water leakage. Carbon paper was glued to the gold connective pads on the PCB using 8331S conductive epoxy (MG Chemicals, Japan), which provides good mechanical and electrical connection. Cathode was prevented from drying by placing an additional layer of porous sponge-based material on the air side of the cathode. Well plate body was 3D printed by Objet 500 Connex 3 3D printer (Stratasys, USA) using compatible Vero resins. Seal was manufactured from PDMS Sylgard 184 (Sigma Aldrich) using a custom-made mold.

**Figure 1.**
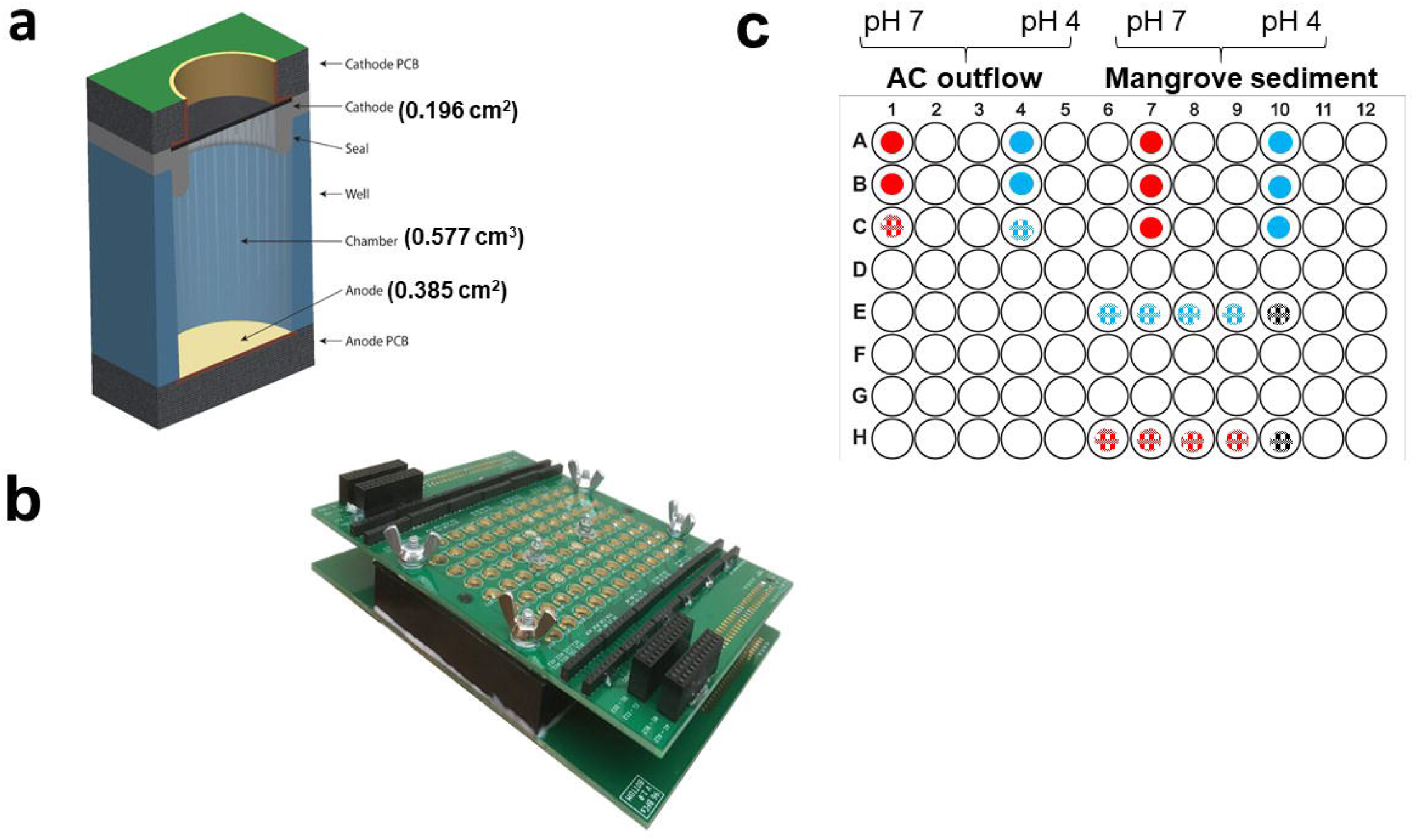
a) Cross-sectional view of the 96 MFCs b) Assembled device c) overview of the experimental setup.

### 2.2. MFC operation

Samples from Okinawa mangrove forests (M), as described in Szydlowski et al. ([16]), and air-conditioning unit outflow in Ishikawa, Japan (26.43°N, 127.84°E; AC) were mixed (1:3) with basal medium containing 200 mg/L CaCl_2_•2H_2_O, 250 mg/L MgCl_2_•6H_2_O, 500 mg/L NH_4_Cl, trace elements and vitamins solution (medium 141 DSMZ) with 1g/L acetate and 2 g/L lactate as carbon source was used, with pH adjusted to either 4 or 7, and incubated overnight at 25°C. To inoculate 96-well plate, five carbon veil (7 gsm, Elite Motoring, USA) disks (0.385 cm^2^ each) were immersed in each solution overnight and transferred to the 96-well plate as indicated in Fig.1c. Disks were placed in wells with R_ext_ = 330 Ohms, as determined by electrochemical impedance spectroscopy (EIS) measurements of the wells (Table S1). Then, the best performing wells from each pH regime were selected, as indicated in Fig. 2a and multiplicated as follows: disk from C1 were placed in wells E6-E10 and disks from were placed into wells H6-H10, wells E10 and H10 serving as open circuit potential (OCP) controls. Additional 4 disks were added to each well to allow biofilm growth. The same medium was used throughout experiment, and the 96-well plate was incubated on bench at 25°C, with fresh medium replaced daily by opening the top PCB and replacing the same volume (577 μl) with pipette. Samples were subjected to VFA analysis using ionchromatography. Top carbon sponges on cathodes were also washed daily with miliQ water. Potential between electrodes with external resistors attached was measured manually, and current was derived using Ohm’s law. Current density (j) was normalized to the volume of each well (0.577 cm^3^) or anode surface area (0.385 cm^2^). After 3 days, the wells with the highest potential were selected and further multiplicated by transferring carbon veil disks into four new wells, with one well serving as OCP control. Potential between electrodes external resistors attached was measured for four weeks using PalmSens3.

**Figure 2.**
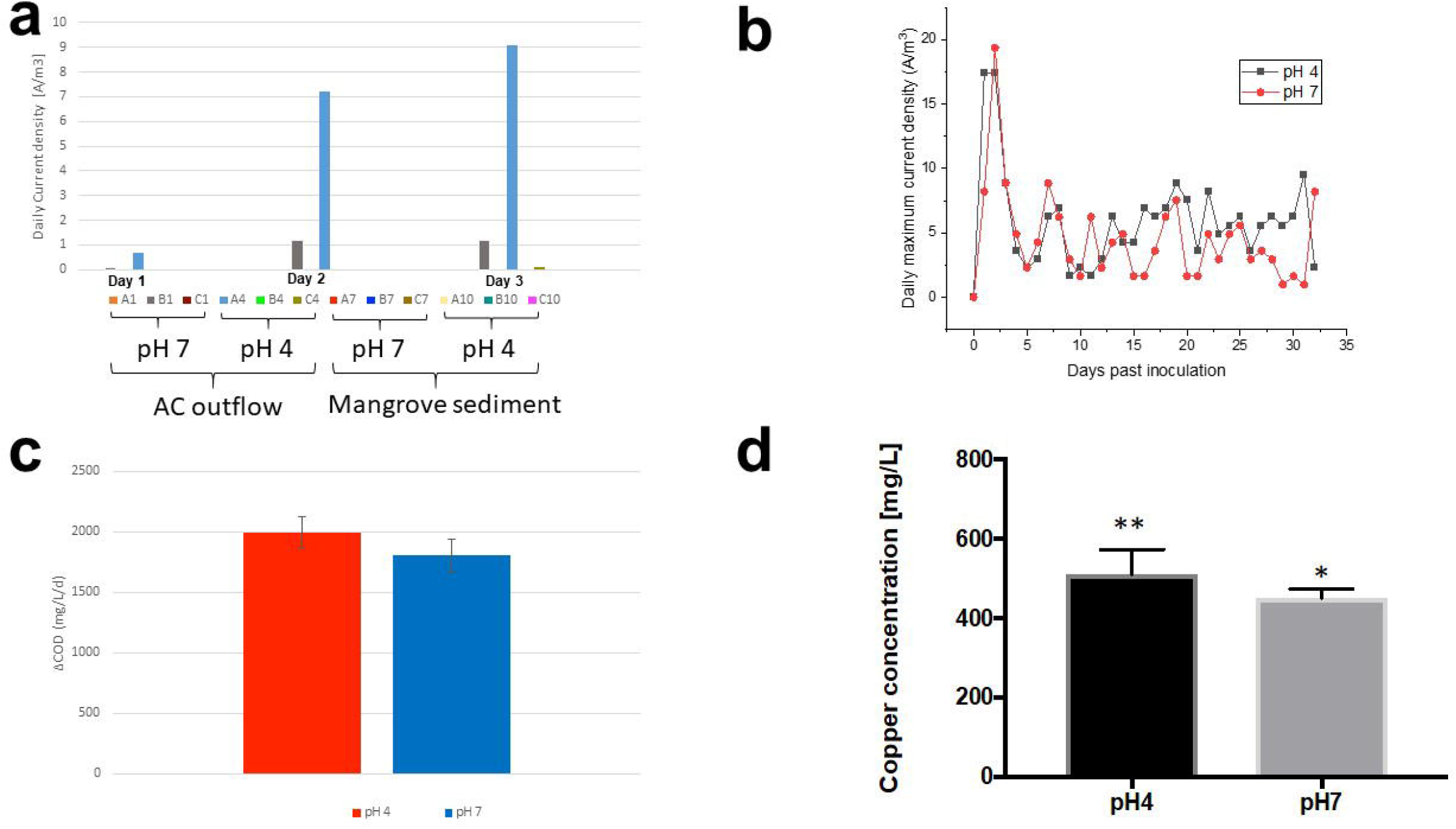
a) Current density measured in all samples 3 days past inoculation, b) daily maximum current density in the best performing wells (E6-9, H6-9), c) COD consumption in the best performing wells (E6, H8), d) copper concentration in the best performing wells (E6, H8).

### 2.3. Electrochemical measurements

Cyclic voltammetry (CV) and linear sweep voltammetry (LSV) were performed using PalmSens3. Electrochemical impedance spectroscopy (EIS) was performed using Gamry Interface 1000. Two-electrode setup was used for measurements after 30 min left at open circuit potential (OCP), with anodes being working electrodes and cathodes being counter and reference electrodes. For CV, following parameters were applied: E range was from 0.4 V to −0.7 V, 0.1 mV/s scan rate. For LSV, the potential range was from 0 mV vs. open circuit potential (OCP) to 0 mV vs. reference electrode, 0.1 mV/s scan rate, and 0.2 mV step. For EIS, the frequency range was between 1 MHz to 50 mHz, with 7 mV steps. EIS data (Table S1) were used to determine the internal resistance of the MFCs.

### 2.4. Copper measurement

Copper concentration from liquid samples was determined using ICP-MS. Samples were diluted 100,000 x with MiliQ water and treated overnight with 5% HNO_3_ to remove residual organic matter.

### 2.5. SEM and EDS observations

Anodes were fixed by soaking in 2.5% glutaraldehyde for 12 hours at 4°C. They were then washed 3 times with 0.1 M phosphate buffer of pH 7 at 4°C, dehydrated with a series of ethanol solutions (50, 70, 80, 90, 95 and three times in 100%). Next, anodes were soaked in pure HMDS twice for 30 seconds, as described elsewhere[17]. After 10 minutes of drying, they were sputter coated with gold. Sample were observed by SEM (JSM-7900F JEOL). Additionally, EDS scans were conducted to detect the presence of copper on biofilms.

### 2.6. DNA sequencing and metagenomic analysis

DNA was extracted using Maxwell RSC kit and automated station. Extracted DNA was subjected to Illumina NovaSeq sequencing and metagenomes were processed using KBase platform [18]. First, paired-end reads were subjected to the QC and filtering using FastQC v0.11.5. For phylogenetic analysis, paired-end libraries were subjected to Kaiju pipeline[19]. Subsequently, metagenmic reads were assembled using metaSPAdes v3.13.0[20]. For functional analysis, contigs were binned using MaxBin2 v2.2.4[21], and extracted genomes quality was assessed with CheckM v1.0.18[22]. Multiple assemblies were annotated using RASTtk[23] and multiple domain annotation tools and subjected to comparative studies.

## 3. Results

### 3.1. MFC operation and electrochemical measurements

During the initial phase, we aimed to observe the startup time for the electrogenic microorganisms, indicated by current generation. After 3 days past inoculation, current reached 5.6 and 0.69 μA in AC outflow samples, which is equivalent to 0.13 and 0.018 A/m^2^ or 9.1 and 1.2 A/m^3^, when calculated against electrode surface area or reactor volume, respectively. No current was observed in mangrove samples (Fig. 2a). We therefore decided to multiplicate communities from wells B1 (pH 7) and A4 (pH 4) derived from AC outflow and connected them to the datalaogger for continuous data collection (Fig. S1). In the first 2 days, current densities reached maximum currents of 17.4 and 19.4 A/m^3^ in pH 4 (well E6) and pH 7 (well H8) samples, respectively. Then, currents decreased and kept oscillating between 1.6 and 8.8 A/m^3^ in the following days, with pH4 wells showing slightly higher values than pH 7 (Fig. 2b). In terms of power, our wells produced up to 30.1 and 80.2 nW (which is equivalent to 0.80 and 0.98 mW/m^2^ or 112.9 and 145.2 mW/m^3^) in E6 (pH 4) and H8 (pH 7) wells, respectively (Fig. S1). Based on the IC analysis, the total COD removal was 1996 ± 127 mg COD/L/d and 1806 ± 137 mg COD/L/d for pH 4 and pH 7 samples, respectively (Fig. 2c). It is also noteworthy that pH had been increasing from 4 to 7 and from 7 to 10 after 24 hours in all samples.

CV scans in pH 4 blank reveals smooth curve non-Faradaic current of ^~^ 30 μA and small reduction peak at +0.55V. Two weeks past inoculation, an oxidative peak is seen at 0.13 V, with oxidative wave at 180 μA and reductive peak at −0.44 V with reduction wave below −200 μA (Fig.3). Therefore, capacitance increase is over 12-fold (from 30 to 380 μA). In pH 7 samples, non-Faradaic currents are below 30 μA, with no redox peaks. Two weeks after inoculation, oxidation peak around 0 V, with oxidation wave at 80 μA and reduction peak −0.25 V with reduction wave at −170 μA are observed (Fig.3). Thus, the capacitance increase is over 8-fold.

**Figure 3.**
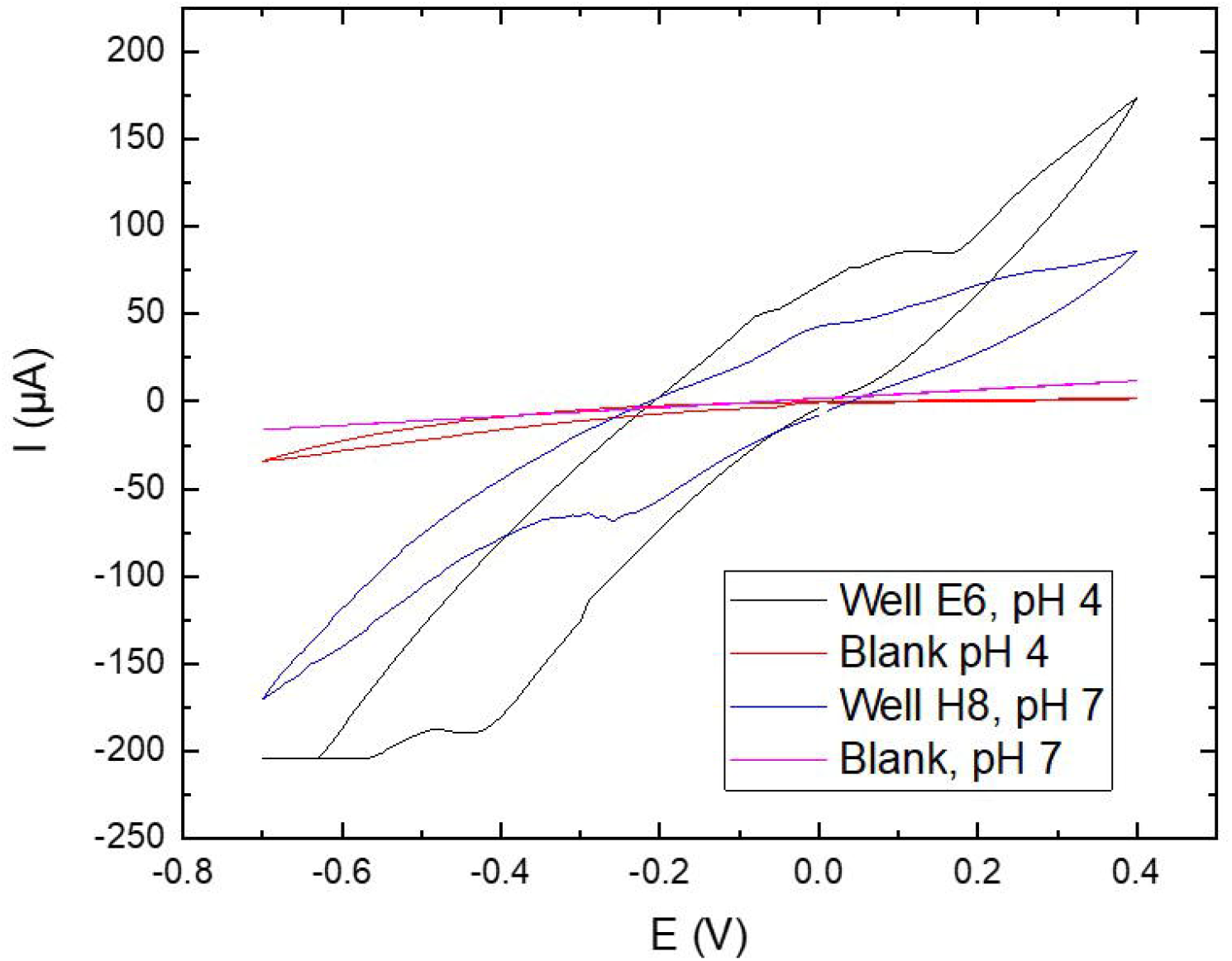
Cyclic voltammetry of blanks and inoculated wells from pH 4 and pH 7 AC samples with blanks.

### 3.2. Copper presence and ICP-MS measurements

After 5 days past initial inoculation, we also noticed blue colour appearing in pH 4 samples anolyte outflow, which we identified as copper ions, potentially derived from PCB. We also noticed greenish salt sediments on the cathode sponges of all samples. We have subjected outflow samples to ICP-MS to determine the concentration of copper was on average 510±63 and 450±24 mg/L for pH 4 and pH 7 samples, respectively (Fig. 2d), whereas the Cu concentration in prepared media was below quantification limit (3 μg/L).

### 3.3. SEM and EDS analysis

The SEM images of anodic biofilms clearly indicate particles adsorbed on cell surfaces (Fig. 4, ab). When further analysis was done using Energy Dispersive Spectroscopy (EDS) detector, strong signal was obtained from copper (Fig. 4 cd). Moreover, in some parts of the biofilm (Fig. 4cd) Cu was present only on some bacterial species, thus suggesting that the community comprises of members actively sequestering Cu ions from the solution.

**Figure 4.**
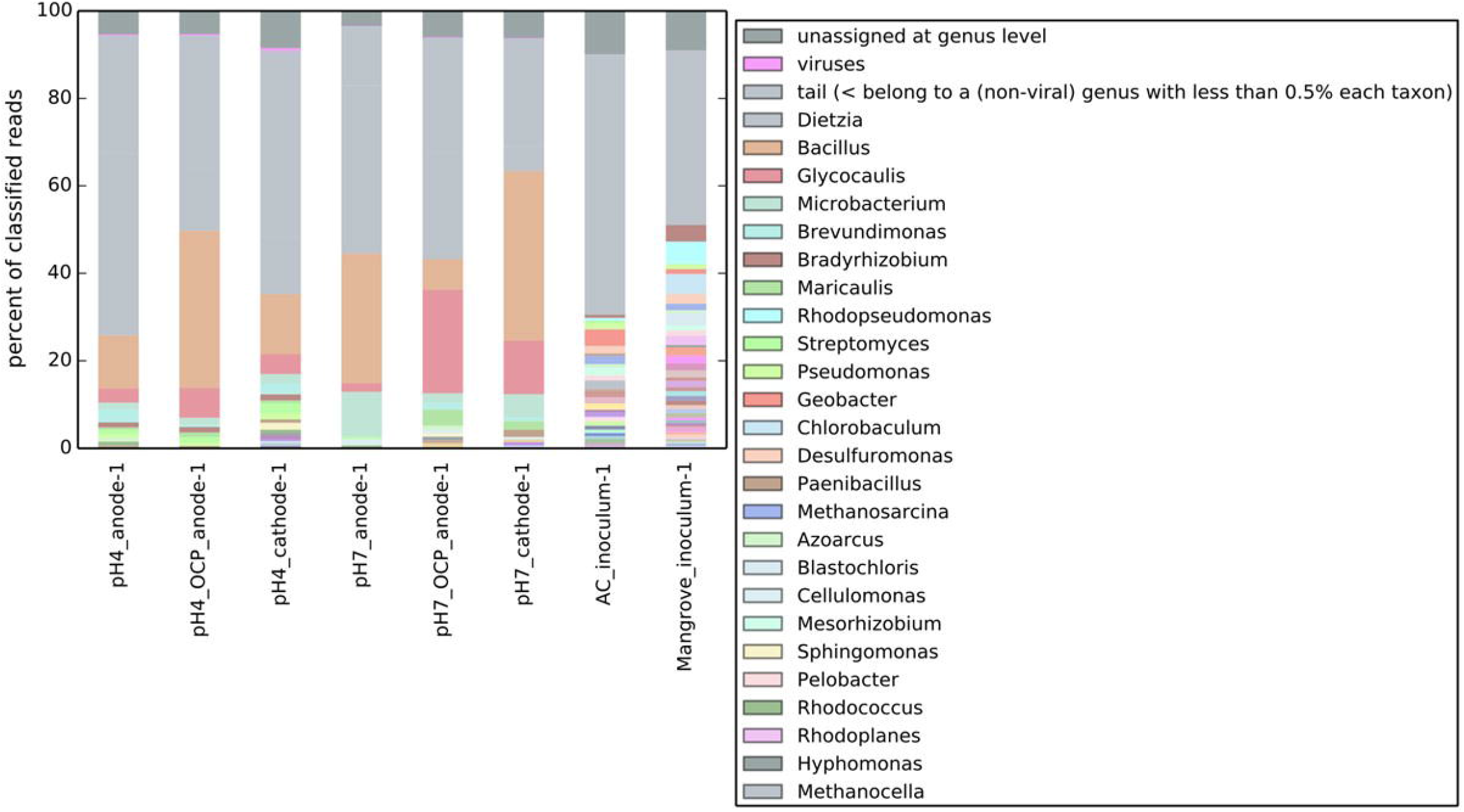
Anodic (a) and OCP (b) biofilm imaging done by SEM and elemental mapping by EDS (c,d).

### 3.4. Metagenomic analysis

Kaiju profiles of our anodic metagenomes reveal that *Dietzia* spp, was the most dominant anodic genus in both pH 4 and 7 reactors derived from AC inoculum. Other abundant genera detected in the samples are *Bacillus, Glycocaulis* and *Microbacterium* that are also abundant in OCP controls. In pH 4 and 7 OCP controls, the most abundant genera are *Bacillus* and *Dietzia/Glycocaulis*, respectively, whereas cathodes comprised mostly of *Bacillus* in both pH groups. Both AC outflow and mangrove sediment, contained *Geobacter*, which was had 4 and 2% abundance, respectively. In AC outflow, *Geobacter* spp. was the most abundant genus, whereas in mangrove sediment, *Chlorobiales* and *Rhodopseudomonas* reached 5% abundance. *Dietzia* comprised only 0.04% and 0.002% of the AC outflow and Mangrove sediment, respectively.

Binned contigs from all pH anodes and OCP controls, revealed genomes of organisms related to *Actinotalea ferriarae, Dietzia psychralkaliphila, Oceanicaulis, Microbacterium, Bacillus*. All aforementioned organisms were found in all anodes and OCP controls except A. *ferriarae*, which was found only in pH 7 anode and its abundance in initial inoculum (AC outflow) was less than 0.002%.

All binned contigs were then subjected to domain annotation using RAST. Comparative analysis revealed the presence of over 523, 6, 9 and 10 unique domains in pH 7 anodes, pH 7 OCP controls, pH 4 anodes and pH 4 OCP controls with 13449 domains being common for all of the groups. Both anodes shared only 3 domains, whereas no domains were shared exclusively between OCP controls (Fig. 6).

**Figure 5.**
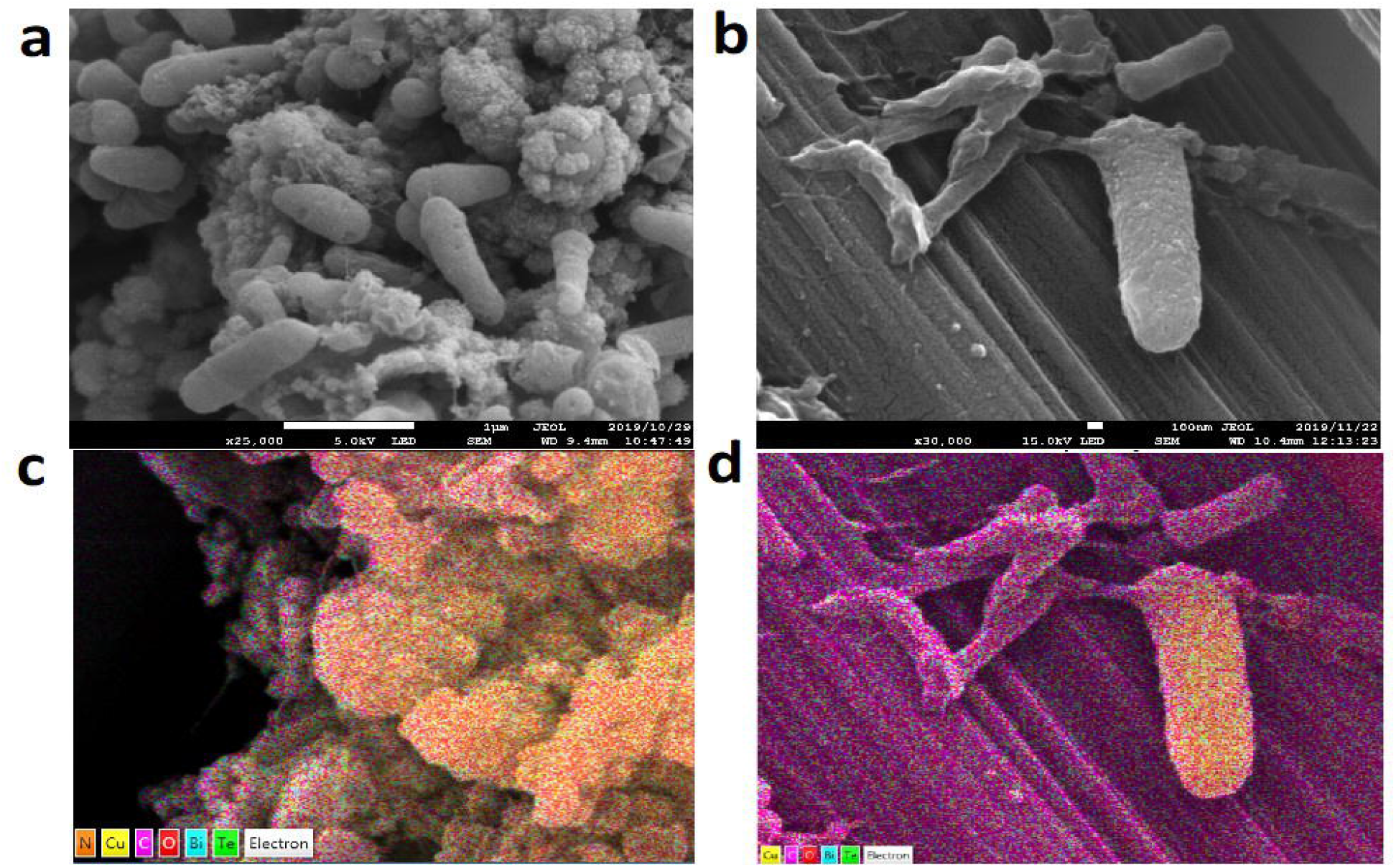
Taxonomy profiles (genus level) of metagenomes using Kaiju. Table shows 25 most abundant genera.

**Figure 6.**
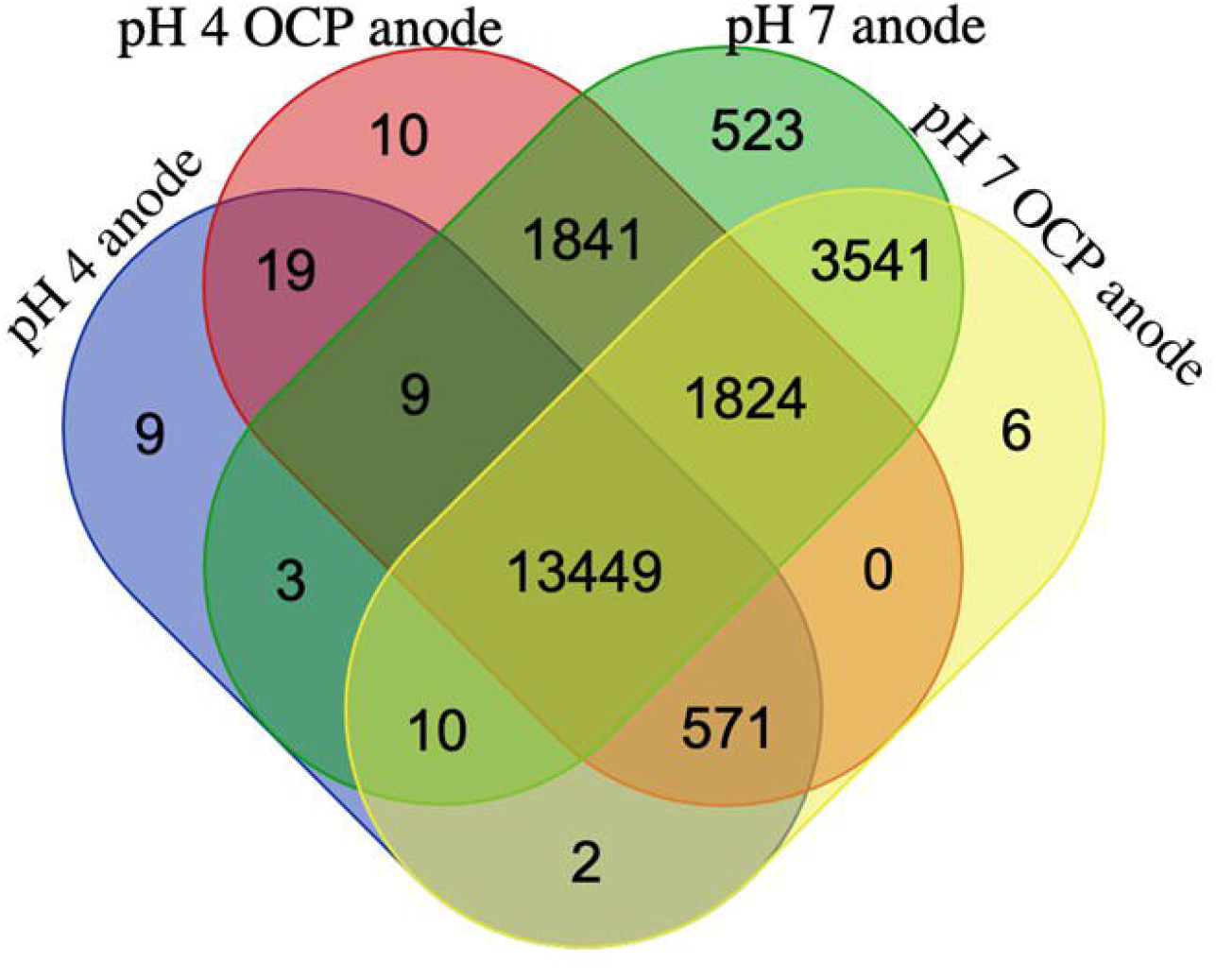
Venn diagram of annotated domains from binned metagenomes. Numbers represent domain annotations found in binned metagenomes from each condition set.

Among unique genes, pH 4 anode bin contain potassium-transporting ATPase A subunit, domains of unknown function. Among OCP control from pH 4, gene encoding vitamin K epoxide reductase (VKOR) is of particular interest. VKOR, a bacterial-specific, membrane protein catalyzes the reduction of vitamin K 2,3-epoxide and vitamin K to vitamin K hydroquinone. It can be also fused with thioredoxin family of oxidoreductases which may function as redox partner to initiate reduction cascade. In pH 7 OCP control, one of the unique annotations is the gene encoding proton-pumping rhodopsin, a photoreactive protein. Anode from pH 7 group has the highest number of unique annotations (523). Among these annotations, 473 originate from the binned genome closely related to *Actinotalea ferriarae*, which is not found in other samples. Within this group, several TRX-like ferredoxin, Ni-Fe and Fe-only hydrogenases, and PQQ enzyme repeats were identified. Other unique annotations, (19 and 15) originate from binned contigs of *Microbacterium* and *Oceanicaulis* spp., respectively.

All binned contigs share genes encoding carbon monoxide dehydrogenase subunit G, as well as metabolic pathways for degradation of various hydrocarbons (both aromatic and aliphatic) and steroids. All contigs also contained various Cu-related genes, such as multicopper- and heme-copper oxidoreductases, as well as genes indicating Cu resistance, e.g., cupredoxins, Copper-resistance genes (*copA*/C/Z), metallochelatins and ATP-dependent heavy metal translocases.

The entire annotation data is available in Supplementary.

## 4. Discussion

### 4.1. Discovery of novel electrogenic species (*Actinotalea* spp.), as well as formerly identified electrogen (Dietzia spp.) that can degrade various hydrocarbons

Our findings reveal that novel electrogenic bacterium has been identified in anodic community. Originally isolated in iron mine [24], this gram-positive, facultatively anaerobic bacterium can survive across wide pH spectrum (4-11), although the most optimal conditions for its growth were in our pH 7 group. *Actinotalea* genus, has been derived from *Cellulomonas*, the main taxonomic difference being the major respiratory quinone: menaquione MK-10(H4) in the former, and MK-9(H3) in the latter [25,26]. Other species from this genus were found in soil [27,28], as well as in biofilm reactor [26]. Presence of several unique annotations in its genome, related to electron transfer confirms that this organism is capable of respiring through anodes.

AC outflow also consists of marine bacterial species originating in deeper seas. *Dietzia* has been previously enriched in MFCs [29], although it is mostly found in oil-contaminated marine environments [30]. Interestingly, despite its alleged alkaliniphility, the abundance of *Dietzia* is similar in both pH 4 and pH 7 anodic samples (Fig. 5) and binned contigs from both samples reveal 99.9 % sequence similarity between the genomes (Fig. S2). This suggests that *Dietzia* is able to withstand periods of lower pH, as the pH increased from 4 to 7 in our pH 4 group. Moreover, its abundance in anodic biofilms may also be explained by the fact that it is capable of degrading hydrocarbons and its toxic by-products, e.g. toluene, phenols etc. Due to the bioleaching of PCB, not only Cu, but also organic epoxy coating gets oxidized, thus creating compounds toxic to many bacteria. It is therefore not surprising that *Dietzia*, being able to degrade numerous hydrocarbons, is so abundant in anodic consortia. Electrochemically-driven release of potentially toxic compounds may also explain why inoculum from Mangrove forest, despite containing electrogenic taxa (*Geobacteraceae*, but no *Actinotalea* and much less *Dietzia*) did not produce current in the same time. Indeed, our attempts to grow pure *Geobacter* cultures in the well plate remained unsuccessful (data not shown).

### 4.2. Enrichment of Cu-resistant, electrogenic community

Despite our efforts to separate PCB from the electrogenic bacteria, our results revealed that chemically deposited Au on electrodes did not provide sufficient barrier against PCB and ultimately led to its corrosion. Owing to this unexpected leakage of Cu ions, we have successfully enriched Cu-resistant communities using our 96-well platform. Through SEM analysis, we observed that anodic biofilm comprises species that concentrate Cu on their cell surface, leaving other parts of the biofilm relatively Cu-free. It is therefore tempting to suggest that this mechanism can be mediated by direct interspecies extracellular electron transfer (DIEET). Indeed, previous works on metallurgic, Cu-removing MFCs demonstrated that electrogenic consortia are capable of reducing Cu from aqueous solution and precipitate it on electrodes. The efficiency of this process depends on various factors, such as the use of pH separators (membranes), presence/absence of oxygen and initial copper concentration [31–35]).

Based on studies on Cu-bioleaching from PCB waste [36–38], one can conclude that Cu mobilisation from its zero-valent state may be due to the bioelectrochemical cycle. Copper can be usually oxidized with simultaneous reduction of Fe^3+^ to Fe^2+^ and its efficiency depends on the reoxidation of iron back to Fe^3+^. Iron oxidation may be catalyzed by bacteria in the presence of oxygen, as well as alternative electron acceptor, e.g. electrode, which in turn may explain high current densities observed in our reactors. Wu et al. [38] demonstrated the use of bacterial-free supernatant derived from Fe/S-oxidizing bacteria, also known for their electrogenic activity, resulted in complete Cu recovery from wasted PCBs.

### 4.3. Demonstration of well plate functionality in terms of culturing electrogenic microbial species

Using our array platform, we have successfully cultivated electrogenic microbial species and monitored their electrochemical performance. Although our system produced less power than that reported by Tahernia and colleagues [12,15], we did not use any catalyst on cathodes, which allows improvement of our device in the future. By altering the conditions between groups (pH), we could distinguish between electrochemical profiles of enriched consortia. However, leaking copper increased conductivity of the medium, thereby affecting current densities and CV profiles, as seen in Figs 2–3. Such problems could be avoided by using the PCBs with so called hard gold finish (electrochemically plated by gold) or covering to PCB anode by other conductive material such as stainless-steel sheet, thus protecting the underlying Cu from being oxidized.

The presented well plate can be used as a standalone technique or in conjunction with other selective processes, e.g. flow cytometry, microfluidic-based dielectrophoretic trapping. Mobile conductive elements, like carbon paper, carbon sponge, activated charcoal granules, metal mesh, can be inserted inside to increase the anode surface area in order to collect electrogenic microorganims and to transfer them into new reactors or for other analytical works. Compatibility of 96 well plate array allows this device to be operated by automated pipetting stations. Modularity of this system allows parts to be disassembled and replaced if needed, furthermore the whole device can be sterilized and reused.

## 5. Acknowledgements

This work has been funded by OIST proof-of-concept programme. We would like to thank Dr. Toshio Sasaki (OIST Imagining analysis) for help with SEM and EDS imaging, Dr. Yoshiteru Iinuma (OIST Instrumental analysis) for chemical analysis, and Mayumi Kawamitsu (OIST Sequencing Section) for their help with metagenome sequencing.

