## Supplementary material for "High-throughput screening and selection of PCB-bioelectrocholeaching, electrogenic microbial communities using single chamber microbial fuel cells based on 96-well plate array": Supplementary.docx

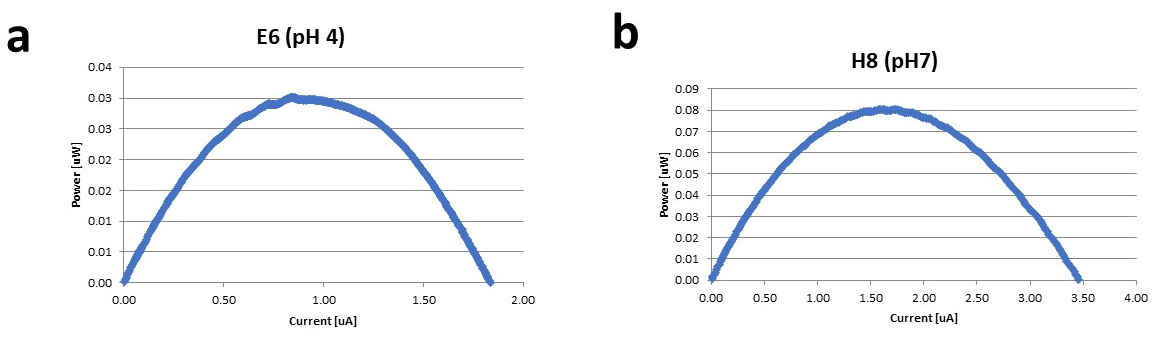

**Figure S1.** Power curves obtained for the best performing wells a) E6 and b) H8.

**Table S1.** EIS measurements of the well plate prior to inoculation. Wells highlighted in yellow were inoculated during this experiment.

|  | 1 | 2 | 3 | 4 | 5 | 6 | 7 | 8 | 9 | 10 | 11 | 12 |
| --- | --- | --- | --- | --- | --- | --- | --- | --- | --- | --- | --- | --- |
| A | 314.6 |  |  | 349.1 |  |  | 325.7 |  |  | 350.3 |  |  |
| B | 337.8 |  |  | 328.9 |  |  | 331.5 |  |  | 310.8 |  |  |
| C | 342.1 |  |  | 310 |  |  | 323.8 |  |  | 345.2 |  |  |
| D |  |  |  |  |  |  |  |  |  |  |  |  |
| E |  |  |  |  |  | 311.9 | 341.6 | 321.6 | 345.4 |  |  |  |
| F |  |  |  |  |  |  |  |  |  |  |  |  |
| G |  |  |  |  |  |  |  |  |  |  |  |  |
| H |  |  |  |  |  | 322.2 | 359.5 | 324.8 | 309 |  |  |  |

**Table S2.** (Separate file). Domains found in annotated metagenomes.
